# Degradation-driven protein level oscillation in the yeast *Saccharomyces cerevisiae*

**DOI:** 10.1101/2021.12.14.472633

**Authors:** Bahareh Mahrou, Azady Pirhanov, Moluk Hadi Alijanvand, Yong Ku Cho, Yong-Jun Shin

## Abstract

Generating robust, predictable perturbations in cellular protein levels will advance our understanding of protein function and enable the control of physiological outcomes in biotechnology applications. Timed periodic changes in protein levels play a critical role in the cell division cycle, cellular stress response, and development. Here we report the generation of robust protein level oscillations by controlling the protein degradation rate in the yeast *Saccharomyces cerevisiae*. Using a photo-sensitive degron and red fluorescent proteins as reporters, we show that under constitutive transcriptional induction, repeated triangular protein level oscillations as fast as 5-10 minute-scale can be generated by modulating the protein degradation rate. Consistent with oscillations generated though transcriptional control, we observed a continuous decrease in the magnitude of oscillations as the input modulation frequency increased, indicating low-pass filtering of input perturbation. By using two red fluorescent proteins with distinct maturation times, we show that the oscillations in protein level is largely unaffected by delays originating from functional protein formation. Our study demonstrates the potential for repeated control of protein levels by controlling the protein degradation rate without altering the transcription rate.

## Introduction

Extensive research has led to insights into how the dynamics and complexity of intracellular regulatory networks are organized to maintain a cellular response (Huang and Ingber 2000, Goldberg 2007, Huang and Ingber 2007, Vogel and Marcotte 2012, Ciechanover 2017). One form of regulation is protein turnover, which refers to the continual renewal or replacement of intracellular proteins. The output of this process is the steady-state intracellular protein concentration, maintained dynamically by continuous production and degradation (Hausser, Mayo et al. 2019). The steady-state intracellular protein level shifts in response to extracellular stimuli or pathophysiologic conditions, which cause changes in production or degradation rates (Baracos 2000, Pratt, Petty et al. 2002, Mizushima and Klionsky 2007, Hinkson and Elias 2011, Ciechanover 2017, Kneppers, Langen et al. 2017).

Interestingly, oscillations in the levels of key proteins have been observed in many biological phenomena (Novák and Tyson 2008), including the cell division cycle (Evans, Rosenthal et al. 1983, Glotzer, Murray et al. 1991), circadian rhythm (Kume, Zylka et al. 1999, Reppert and Weaver 2001), cellular stress response (Lahav, Rosenfeld et al. 2004, Geva-Zatorsky, Rosenfeld et al. 2006), and development (Hirata, Yoshiura et al. 2002, Pourquié 2003). In cells, the oscillation of protein levels is achieved by coordinating of many factors, such as transcriptional activities, translation, post-translational control (e.g., phosphorylation), nuclear transport, and targeted protein degradation. A common feature in oscillations is the feedback loop, which can be either positive or negative (Goldbeter 2002, Novák and Tyson 2008). Theoretical models using negative feedback with a time delay can explain a wide range of protein level oscillations (Lev Bar-Or, Maya et al. 2000, Monk 2003, Tiana, Krishna et al. 2007, Novák and Tyson 2008).

It has been hypothesized that such oscillations enable cells to adapt to periodic changes in the environment (Dunlap 1999), work as a low-pass filter to withstand high-frequency oscillations (Bennett, Pang et al. 2008), repeated repair following DNA damage (Batchelor, Loewer et al. 2009), generate multiple gene expression patterns (Wei, Wu et al. 2006), and potentially integrate multiple cycles of signals before apoptosis (Lahav 2004). A means to elucidate the causal role of protein level oscillations in a key protein would be a robust method to generate timed protein level changes using orthogonal synthetic control decoupled from endogenous signals.

Many versions of synthetic genetic oscillators have been realized through transcriptional control (Novák and Tyson 2008, Purcell, Savery et al. 2010). Although these synthetic design and implementation of oscillators in cells has played a critical role in elucidating the origin and robustness of oscillations (Vilar, Kueh et al. 2002, Purcell, Savery et al. 2010, Woods, Leon et al. 2016), controlling the protein degradation rate has not been explored in synthetic protein-level oscillations. In natural systems, a varying rate of targeted protein degradation has been widely observed in oscillations (Glotzer, Murray et al. 1991, Varshavsky 2008, Batchelor, Loewer et al. 2009). A prominent example is the oscillatory dynamics observed in the tumor suppressor protein p53. Upon DNA damage by γ radiation, the p53 level oscillates without dampening for days (Geva□atorsky, Rosenfeld et al. 2006) with the number of pulses increasing as the irradiation dose increases (Lahav, Rosenfeld et al. 2004). The oscillation of p53 is driven by a negative feedback loop between p53 and Mdm2, in which p53 transcriptionally activates Mdm2, which then binds to p53 and mediates its degradation (Piette, Neel et al. 1997, Lakin and Jackson 1999, Batchelor, Loewer et al. 2009).

Various experimental tools for controlling protein levels have been developed. An extensive list of regulators is available at the transcription level, many of which have been implemented for a wide range of applications (Tigges and Fussenegger 2009, Engstrom and Pfleger 2017, Ding, Zhou et al. 2021). The protein degradation rate can be modulated using degrons, which are short peptide sequences that bind to ubiquitin ligases (Varshavsky 2008, Varshavsky 2019). Harnessing conditional degrons (Natsume and Kanemaki 2017, Luh, Scheib et al. 2020), the protein degradation rate can be controlled by temperature change (Dohmen, Wu et al. 1994, Faden, Ramezani et al. 2016), small molecule binding (Stankunas, Bayle et al. 2003, Banaszynski, Chen et al. 2006, Nishimura, Fukagawa et al. 2009), protease cleavage (Taxis, Stier et al. 2009), or light exposure (Renicke, Schuster et al. 2013, Bonger, Rakhit et al. 2014).

We applied a photo-sensitive degron (psd) (Renicke, Schuster et al. 2013) in *S. cerevisiae* to study the dynamics of protein level oscillations driven by changes in the degradation rate. In this approach, a reporter protein is fused to a photoreceptor protein domain that changes its C-terminal conformation upon blue light exposure. Under blue light, the C-terminally fused degron becomes more accessible, resulting in enhanced protein degradation (**Fig. 1**). Photo-activation provides unique advantages for modulating protein turnover due to the ease of temporal and spatial control of light delivery. In addition, many cell types are not sensitive to light, making them suitable for orthogonal control. We took advantage of the psd developed by Renicke and colleagues to generate frequency-dependent oscillations in protein levels, induced by varying frequency of light inputs to modulate the protein degradation.

**Figure 1.**
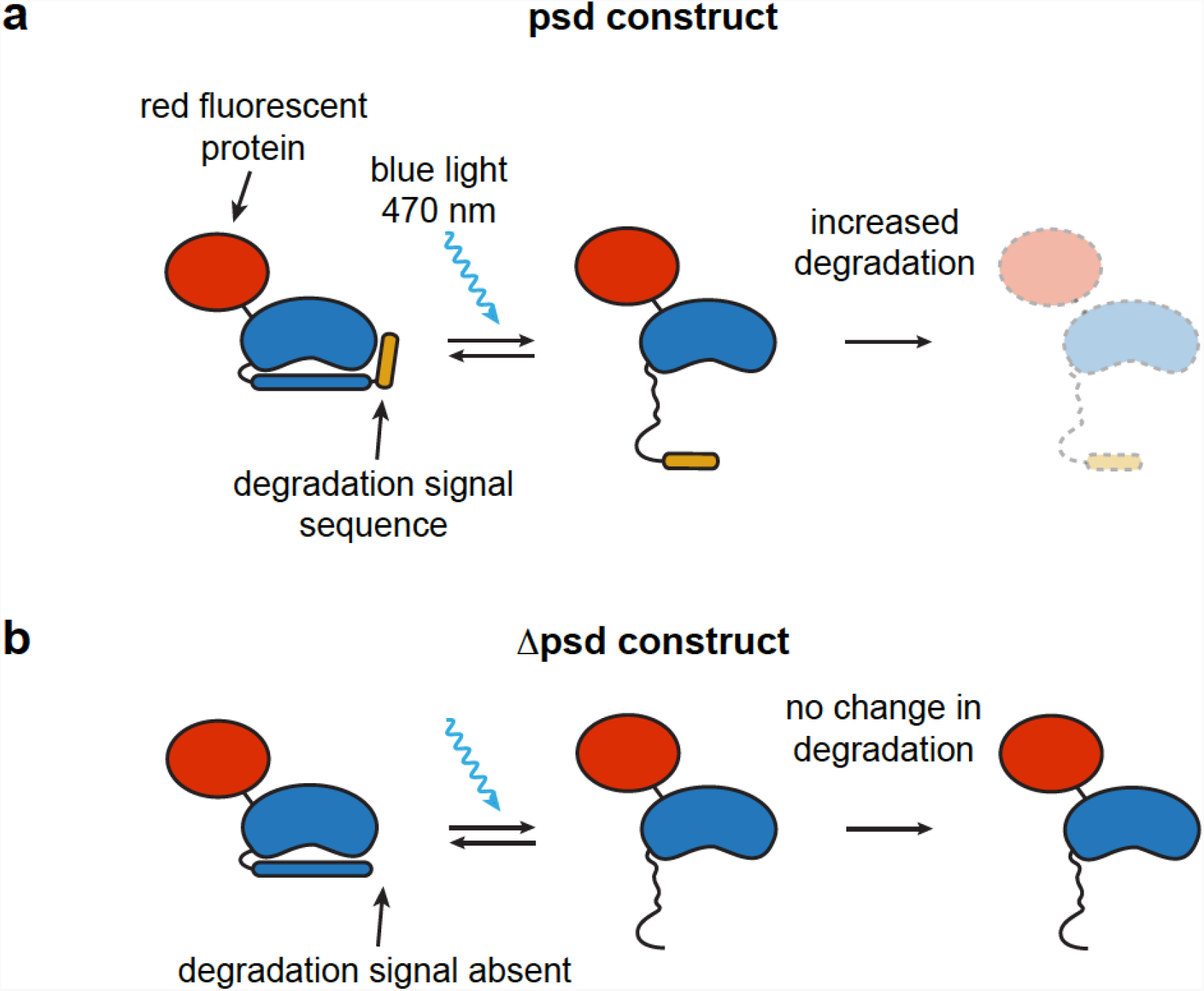
Schematic of photo-sensitive degron (psd) constructs used in this study. In the psd construct (**a**), a protein degradation signal sequence derived from the ornithine decarboxylase is fused to the blue-light sensing light-oxygen-voltage (LOV) domain from *Arabidopsis thaliana*. Upon blue light activation, the degradation signal accessibility increases, resulting in faster protein degradation. The Δpsd construct (**b**), contains all components of psd except the degradation signal sequence.

Using this system, we observed that robust protein level oscillations can be generated by modulating only the protein degradation rate under constitutive induction. The frequency of oscillation followed that of the input light illumination frequency down to the 5-10 minute scale. We also compared the degradation-driven oscillations of two red fluorescent proteins with distinct maturation times to assess how delays originating during functional protein formation affect protein level oscillation. By quantitatively comparing the oscillation amplitude and phase shift as a function of input frequency, we demonstrate that the oscillations at the protein level are largely unaffected by production delays. Overall, we find that protein level oscillations can be robustly generated simply by modulating the protein degradation rate.

## Results and Discussion

### Degradation-driven protein level oscillations in the yeast *S. cerevisiae*

The psd module (**Fig. 1**) comprises the light oxygen voltage 2 domain from *Arabidopsis thaliana* (*At*LOV2), fused with a derivative of the carboxyl-terminal degron of murine ornithine decarboxylase (ODC), enabling the control of the ubiquitin-independent proteasomal degradation rate on blue light illumination (Renicke, Schuster et al. 2013). While *At*LOV2 is sensitive and responsive to blue light, the ODC degradation signal is responsible for increased protein degradation (**Fig. 1**). To observe the effect of modulating the protein degradation rate on protein turnover, we applied pulses of blue light and monitored the fluorescence emission from the red fluorescent protein tagRFP fused to psd (**Fig. 1a**). To validate protein degradation due to the ODC degradation signal, we generated a control construct lacking the signal sequence, termed Δpsd (**Fig. 1b**). The expression of these constructs in yeast *S. cerevisiae* was continuously driven using the GAL_1-10_ promoter under galactose. To validate continuous expression, we measured the *tagRFP* transcript level by extracting the mRNA from yeast cells and performing a quantitative polymerase chain reaction (qPCR) of the transcript (**Fig. 2a**, see **Supp. Fig. 1** and **Supp. Table 1** for RNA quality control and primers used). Previous studies indicated that prolonged exposure (> 10 hr) of extremely high intensity light (> 10-fold higher intensity used in this study) causes oxidative stress in yeast (Bodvard, Wrangborg et al. 2011, Robertson, Davis et al. 2013). To assess any potential effect of illumination on gene expression, we illuminated the cells with the maximum blue light intensity used in this study (light power density of 0.027 mW/mm^2^ [106.02 µmol/m^2^s]) and the longest duration (1/*f* = 100 min where *f* indicates the illumination frequency). The mRNA was extracted from cells at the start of the experiment and at the end of each illumination cycle (see inset in **Fig. 2a**). Two primer sets binding to different regions of *tagRFP* were used for amplification, along with primers for two previously validated housekeeping genes, *TFC1* and *UBC6* (Teste, Duquenne et al. 2009), as a reference. The expression of *tagRFP* remained unchanged for 200 min, and a significant increase was detected between 200 and 300 min (**Fig. 2a**, *p =* 0.014 and 0.023, student’s t-test comparing ΔCt at 200 min. to that at 300 min. for the two *tagRFP* primer sets, respectively [n = 3 independent culture preparations]). This result indicates that the GAL_1-10_ promoter allows continuous expression of the tagRFP constructs, unhindered by light illumination.

**Figure 2.**
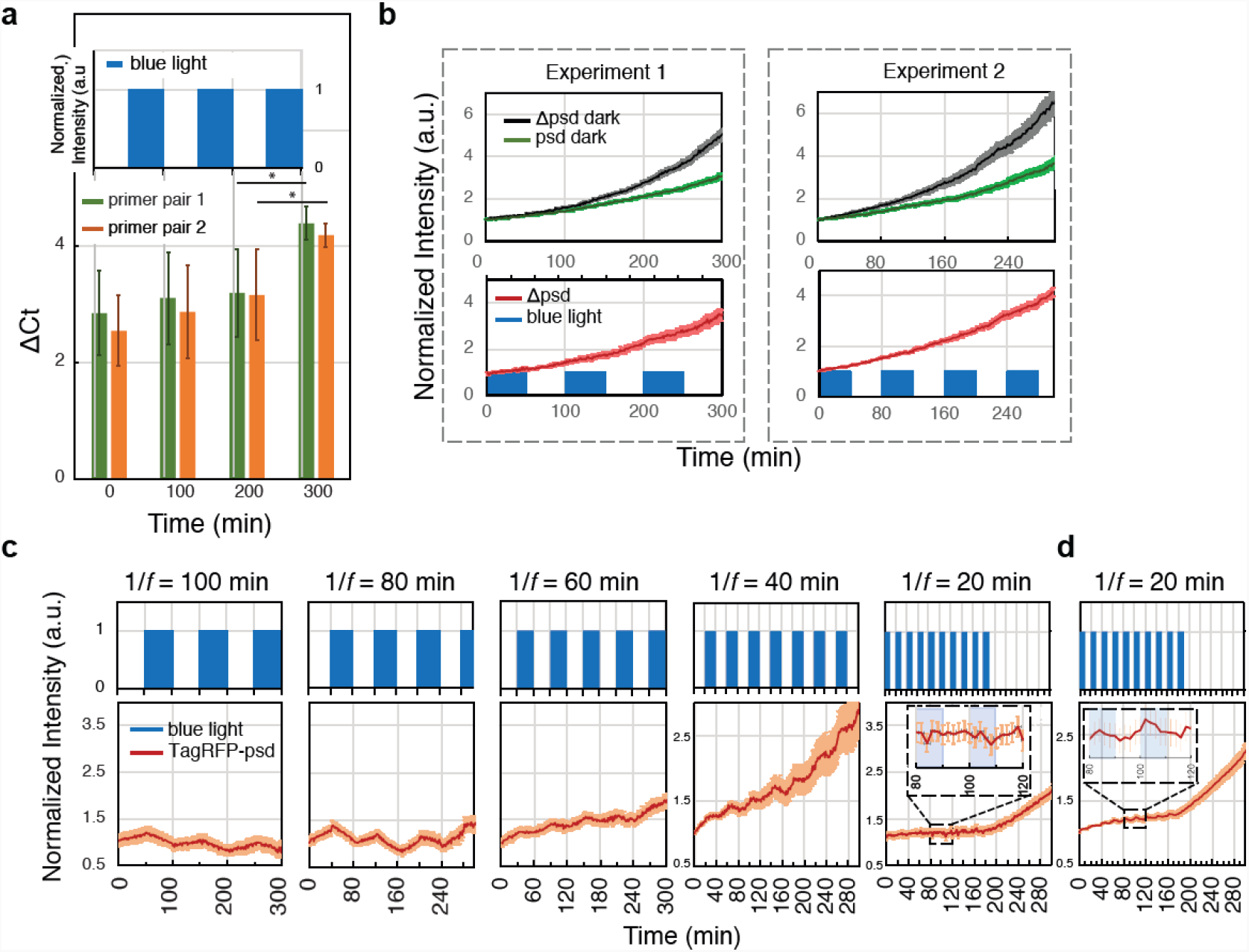
Experimental validation and observation of degradation-driven protein level oscillations in yeast expressing tagRFP fused to psd or Δpsd. (**a**) qPCR analysis of *tagRFP* transcripts during blue light illumination (inset indicates illumination timepoints). Two sets of primers (primer sets 1 and 2) binding to different regions of *tagRFP* were used, and previously validated housekeeping genes *TFC1* and *UBC6* were used as references. The ΔCt values were calculated by subtracting the Ct of *tagRFP* from the average of the Ct of the two reference genes. *: *p <* 0.05, student’s t-test comparing ΔCt at 200 min. to that at 300 min. Error bars indicate standard deviation from n = 3 independent culture preparations. (**b**) Normalized fluorescence levels of tagRFP-psd and tagRFP-Δpsd in the dark (top panels); tagRFP-Δpsd illuminated with blue light cycles (bottom panels). Solid lines indicate the mean fluorescence intensities and lighter lines indicate the standard deviation from n = 10 to 15 microcolonies. (**c**) Fluorescence of tagRFP-psd under varying frequencies of blue light illumination. *f* denotes the temporal frequency and 1/*f* indicates the cycle period. Solid lines indicate the mean fluorescence intensities and lighter lines indicate standard deviation from n = 10 to 15 microcolonies. (**d**) Experiment for 1/*f* = 20 min. was repeated with 25 microcolonies. The same intensity of blue light (power density of 0.027 mW/mm^2^ [106.02 µmol/m^2^s]) was used in all panels. In panels (b) and (c), data for each illumination frequency were obtained in separate experiments using independent cell cultures and each microcolony contained 4 to 8 cells at the beginning of experiment.

Since the cell cycle of *S. cerevisiae* is of the order of 100 min (Herskowitz 1988), we observed cell division during the repeated illumination cycles (**Supp. Fig. 2**). The proteins in each cell are divided into daughter cells, resulting in protein dilution. One approach to quantify all the proteins throughout cell division is to sum the fluorescence intensity from a microcolony of cells originating from mother cells (Talia, Skotheim et al. 2007). Consistent with other studies, we found that summing the fluorescence intensity of a microcolony of cells resulted in a continuous increase (red line in **Supp. Fig. 2**), while the fluorescence intensity divided by the area of cells fluctuated as the cells divided (blue line in **Supp. Fig. 2**). To capture the change in protein level decoupled from cell division, we used the sum of the fluorescence intensities of a microcolony of cells as a measure of protein levels throughout this study.

We then tested the effect of pulses of blue light on yeast cells expressing tagRFP-psd or tagRFP-Δpsd. In the absence of blue light, the normalized fluorescence level of tagRFP-psd and tagRFP-Δpsd (see **Materials and Methods** section for calculation) showed a continuous increase over 300 min. (**Fig. 2b**). The fluorescence level of tagRFP-Δpsd was generally higher than that of tagRFP-psd (**Fig. 2b**), probably because the LOV domain conformation was in equilibrium between active and inactive states even in the dark (Strickland, Yao et al. 2010), resulting in slight background degradation even without blue illumination. As anticipated, the level of tagRFP-Δpsd was unresponsive to two distinct illumination frequencies (**Fig. 2b**, *f* = 1/100 min and *f* = 1/80 min, respectively in experiments 1 and 2). Conversely, the fluorescence level of tagRFP-psd showed distinct oscillations with varying frequencies of blue light (**Fig. 2c**). We repeated the experiment five times, each at a distinct illumination frequency (**Fig. 2c**). Protein level oscillation was clearly visible up to an illumination frequency of 1/*f* = 40 min, but not at the highest illumination frequency tested (1/*f* = 20 min) (**Fig. 2c**). When the experiment for 1/*f* = 20 min was repeated with a larger number of microcolonies, the oscillation was discernable albeit markedly reduced (**Fig. 2d**). Based on these findings, we conclude that the output oscillations observed in the experiment were the result of periodic light-induced alterations in the protein degradation rate mediated by the psd.

### Degradation-driven oscillations in a faster maturing fluorescent protein

Fluorescent proteins contain a chromophore formed through autocatalytic maturation (Wachter 2007, Craggs 2009, Subach and Verkhusha 2012). The fluorescence is only detected after maturation, effectively causing a decrease in the apparent protein production rate. Although not tested, a shift in protein maturation time is expected to impact the low-pass filtering behavior when the transcription rate is modulated as in Bennett *et al*.’s work (Bennett, Pang et al. 2008). Conversely, when the protein degradation rate is modulated, as in our experiments, a shift in protein maturation time should not affect the oscillation behavior. Validating this would indicate that by modulating the protein degradation rate, consistent protein level oscillations can be generated for proteins with distinct production rates.

To test this, we replaced tagRFP in tagRFP-psd with the fluorescent protein mCherry (mCherry-psd), which has a significantly shorter maturation time compared to that of tagRFP (maturation half-time of ∼40 min for mCherry vs. ∼100 min for tagRFP (Merzlyak, Goedhart et al. 2007)). We then repeated the experiment shown in **Fig. 2c** using mCherry-psd and monitored the mCherry fluorescence in microcolony numbers similar to that in **Fig. 2d** (25 to 30 microcolonies of 4 to 8 cells at the beginning of the experiment). Clear oscillations in mCherry fluorescence were visible, down to an illumination frequency of 1/*f* = 20 min (**Fig. 3a**). We also conducted more experiments with higher input frequencies (1/*f* = 10 min and 1/*f* = 5 min) and detected small but distinguishable oscillations (**Fig. 3b** and see **Supp. Fig. 4** for extended experiments). We observed consistent decrease in the amplitudes of output oscillations on increasing the frequency of degradation oscillation, characteristic of a low-pass filtering behavior.

**Figure 3.**
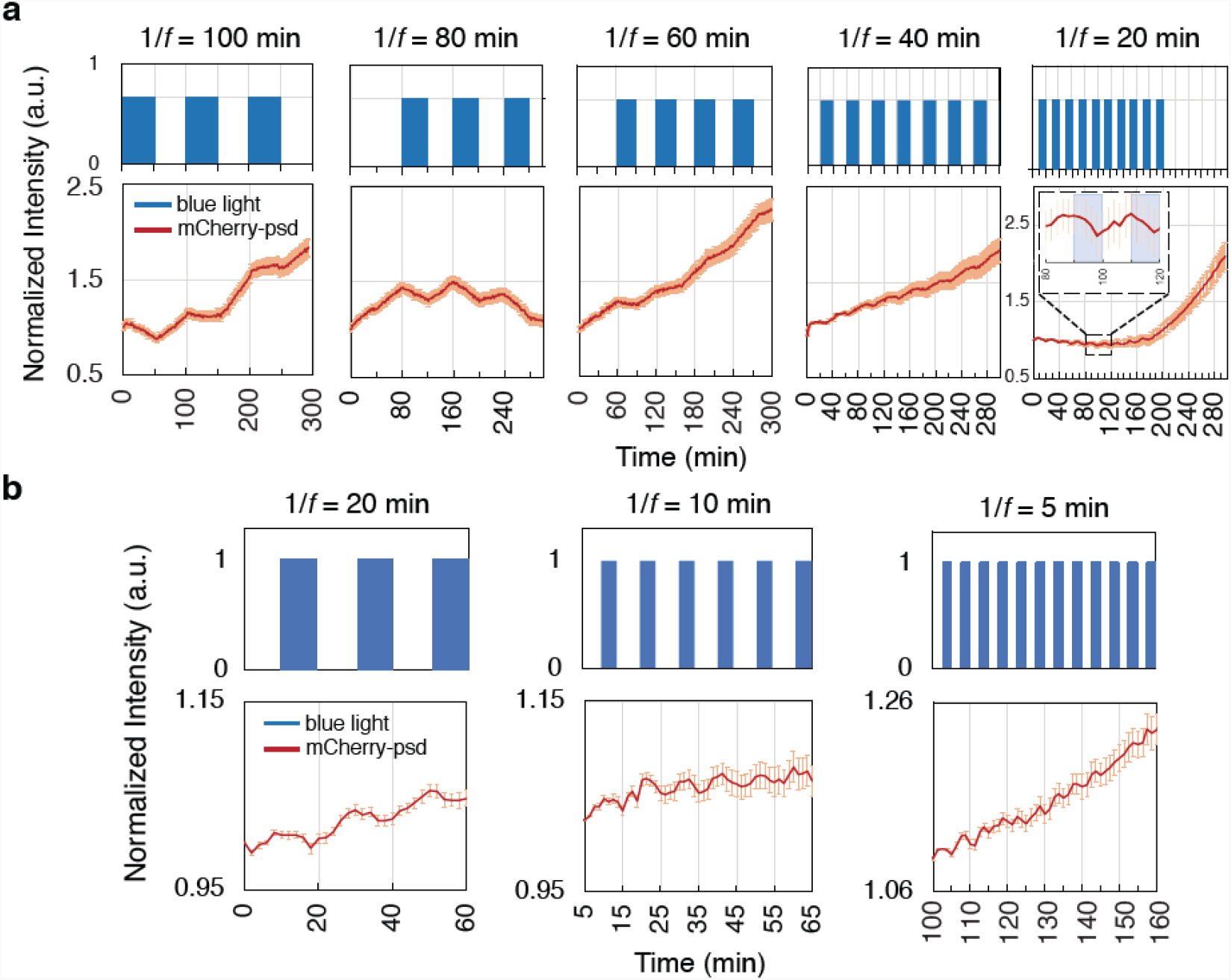
Experimental observation of degradation-driven protein level oscillations in yeast expressing mCherry fused to psd. (**a**) Fluorescence of mCherry-psd under varying frequencies of blue light illumination. (**b**) Fluorescence of mCherry-psd under higher frequency illumination. For frequency 1/*f* = 20 min., the experiment in panel (a) was repeated. For all panels, solid lines indicate the mean fluorescence intensities and lighter lines indicate the standard deviation from n = 25 to 30 microcolonies. The blue light intensity was 0.027 mW/mm^2^ (106.02 µmol/m^2^s)) for all panels. Each microcolony contained 4 to 8 cells at the beginning of experiment.

### System identification and frequency-domain analysis

The reduction in oscillations in protein levels at higher input illumination frequencies is characteristic of low-pass filtering (Antoniou 2016), and indicates that the input perturbation driving protein degradation was filtered out at these higher frequencies. To analyze the low-pass behavior of the degradation driven protein turnover, we determined the time delay between th input blue light illumination frequency and output protein fluorescence intensity using cross-correlation. We calculated the cross-correlation (see **Supplementary Information** for calculation) for experiments using tagRFP-psd and mCherry-psd for frequencies 1/*f* = 40, 60, 80, and 100 (**Fig. 4**). Since the protein level decreased on illumination, a minimum occurred in the cross-correlation (**Fig. 4**, indicated by arrows). Based on the minimum value for each experiment, the time delay between the input and output signals ranged from 1 to 15 min (see **Supplementary Information** for calculation). For both tagRFP-psd and mCherry-psd, we found that the time at which minima occurred shortened as the input frequency oscillations increased, characteristic of a low-pass filtering system.

**Figure 4.**
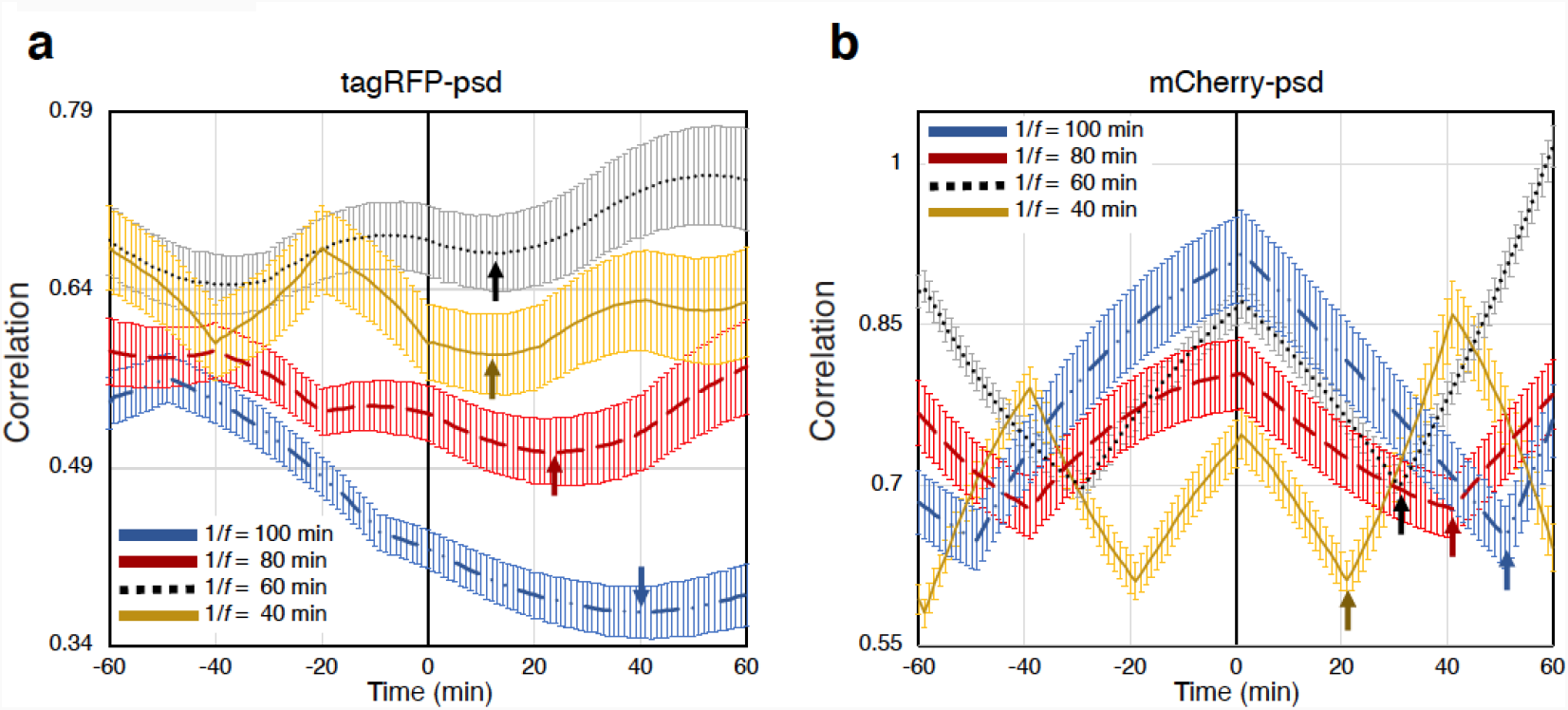
Cross-correlation between input blue light illumination and protein fluorescence intensity. The curves are cross-correlations resulted from experiments with varying input frequencies for tagRFP-psd (**a**) and mCherry-psd (**b**). The arrows show the minimum (after time 0) of each curve. Note that the minima do not directly indicate the time delay but allow the calculation of the time delay as shown in the Supplementary Information.

To quantitatively assess the low-pass filtering, we constructed a mathematical model for degradation-driven protein turnover using the difference equation approach we previously described (Shin 2016). The model equation can be expressed as

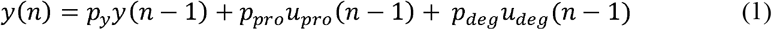

where n is the discrete-time index, y(n) is the normalized fluorescence intensity at time n, u_pro_(n) is the normalized amount of galactose (transcription inducer) at time n, and u_deg_(n) is the normalized intensity of blue light (degradation rate controller) at time n. Parameters p_y_, p_pro_, and p_deg_ indicate endogenous degradation, production, and light-induced degradation, respectively. These parameters can be adaptively estimated from the experimental data using the normalized least mean square (NLMS) algorithm, one of commonly used adaptive system identification algorithm (Ljung 1998, Sayed 2011). Assuming constant protein production, as indicated by relatively consistent transcription (**Fig. 2a**), Equation 1 can be simplified by approximating u_pro_(n) to 1:

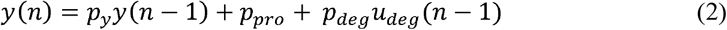

Equation 2 represents a multiple-input single-output (MISO) system that can be transformed into the frequency domain using the z–transform.

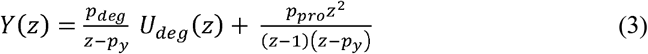

Equation 3 illustrates the relationship between the single output Y(z) and two independent inputs (blue light intensity, U_deg_(z), and the normalized galactose concentration which is assumed to be constant at 1). The equation can be reexpressed in matrix form:

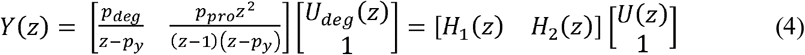

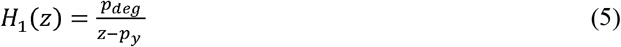

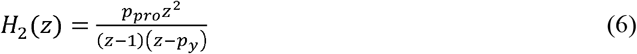

Here, H_1_(z) shows the relationship between the output Y(z) (protein fluorescence intensity) and input U_deg_(z) (blue light intensity). The unknown parameter values p_y_ and p_deg_ in H_1_(z) were estimated using the NLMS adaptive system identification algorithm (See **Supplementary Information** and **Supp. Fig. 5**). The Bode plot in **Fig. 5** (indicated as “Model”) shows the amplitude and phase shift response of H_1_(z), calculated based on the mean and standard deviation of estimated parameter values acquired from each experiment (See **Supplementary Information**). The amplitude responses demonstrate low-pass filtering, indicated by a steady decrease in the oscillation amplitude as the input blue light illumination frequency increases (**Fig. 5a**). We performed a statistical comparison of the experimental results (tagRFP-psd vs. mCherry-psd) and experiments to the model (tagRFP-psd vs. model, mCherry-psd vs. model). The summary of statistical comparisons in **Table 1** shows that the model effectively captured the amplitude and phase shift responses well. Regarding the amplitude response, no significant difference was found when comparing the model vs. tagRFP-psd or the model vs. mCherry-psd (**Fig. 5a** and **Table 1**). When comparing the phase shift response of the model vs. tagRFP-psd, a significant difference was found for 1/*f* = 100 min, 1/*f* = 80 min, and 1/*f* = 60 min but not at other frequencies (**Fig. 5b** and **Table 1**). The phase shift response of the model vs. mCherry-psd was significantly different only at illumination frequency of 1/*f* = 100 min (**Fig. 5b** and **Table 1**). When comparing tagRFP-psd vs. mCherry-psd, mCherry-psd showed significantly smaller amplitude responses at frequencies 1/*f* = 80 min, 1/*f* = 60 min, and 1/*f* = 40 min (**Fig. 5a** and **Table 1**), and significantly smaller phase shift at 1/*f* = 60 min, but no difference was found at other frequencies (**Fig. 5b** and **Table 1**). These results show that both tagRFP-psd and mCherry-psd showed low-pass filtering with similar time delays and confirm that the difference in maturation time between tagRFP and mCherry does not cause a significant change in oscillatory behavior at a wide range of frequencies. Comparing the Bode plot of our experiments with the previous study using transcriptional control (Bennett, Pang et al. 2008), we found that the same amplitude of oscillations (e.g., an amplitude of around 0.3 which is equivalent to -10.5 dB) was achieved at approximately 3-fold higher frequencies (0.06 rad/min) under degradation-driven control (achieved at frequency of 0.02 rad/min), indicating that larger oscillations can be generated at higher frequencies. Taken together, these results demonstrate that degradation driven control is a robust approach to generating protein level oscillations on minute timescales.

**Figure 5.**
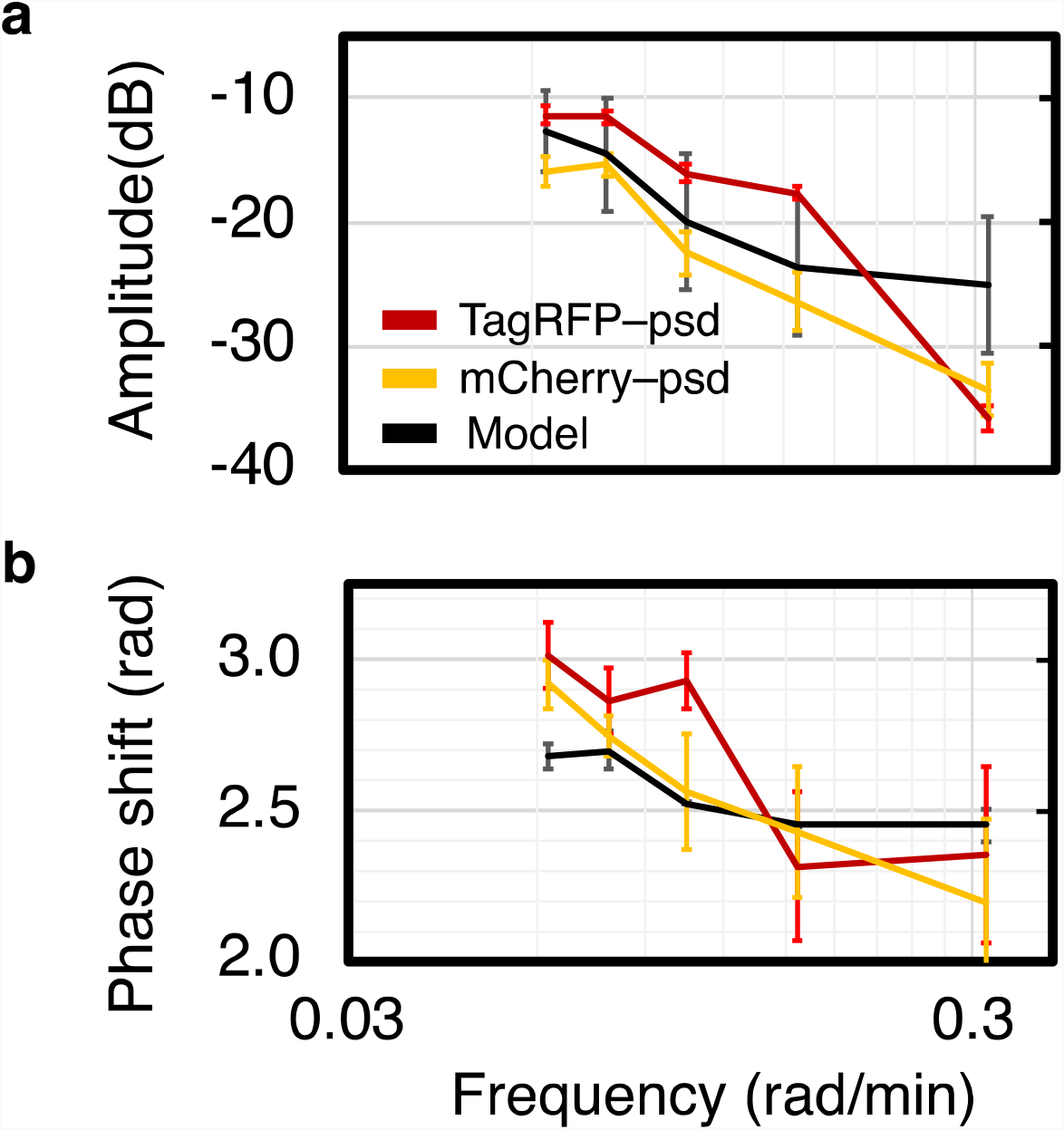
Model and experimental comparison of low-pass filtering using Bode plots. (**a**) Amplitude response and (**b**) phase shift response from tagRFP-psd, mCherry-psd, and model calculations as a function of the input frequency. Data points indicate the median and error bars indicate the interquartile range (IQR) from 10 microcolonies. Statistical analysis of the results is provided in Table 1.

**Table 1.**

| Group \ Set | f = 2π/100<br>(1/f = 100 min) | | f = 2π/80<br>(1/f = 80 min) | | f = 2π/60<br>(1/f = 60 min) | | f = 2π/40<br>(1/f = 40 min) | | f = 2π/20<br>(1/f = 20 min) | |
| --- | --- | --- | --- | --- | --- | --- | --- | --- | --- | --- |
|  | Amplitude | Phase shift | Amplitude | Phase shift | Amplitude | Phase shift | Amplitude | Phase shift | Amplitude | Phase shift |
|  | Median (interquartile range, IQR) |  |  |  |  |  |  |  |  |  |
| Model | -12.691<br>(6.657) | 2.681<br>(0.081) | -14.579<br>(9.136) | 2.701<br>(0.127) | -20.007<br>(10.840) | 2.527<br>(0.008) | -23.572<br>(11.131) | 2.454<br>(0.002) | -25.096<br>(4.847) | 2.454<br>(11.196) |
| tagRFP-psd | -11.373<br>(1.498) | 3.016<br>(0.220) | -11.537<br>(1.149) | 2.867<br>(0.216) | -16.055<br>(1.421) | 2.932<br>(0.183) | -17.657<br>(1.205) | 2.317<br>(0.491) | -35.654<br>(2.146) | 2.356<br>(0.589) |
| mCherry-psd | -15.918<br>(2.385) | 2.922<br>(0.157) | -15.391<br>(1.972) | 2.749<br>(0.137) | -22.518<br>(3.352) | 2.566<br>(0.380) | -26.478<br>(4.660) | 2.435<br>(0.432) | -34.437<br>(4.413) | 2.199<br>(0.550) |
| P-value Model vs tagRFP-psd | 0.438533 | 0.0001 | 0.079751 | 0.0017 | 0.063264 | 0.0066 | 0.079751 | 0.7342 | 0.330421 | 0.9231 |
| P-value Model vs mCherry-psd | 0.620585 | 0.0019 | 0.918717 | 0.1961 | 0.979021 | 0.9541 | 0.918717 | 0.9851 | 0.620585 | 0.4881 |
| P-value tagRFP-psd vs mCherry-psd | 0.0842 | 0.7265 | 0.0094 | 0.1961 | 0.0049 | 0.0161 | 0.0069 | 0.8287 | 0.5037 | 0.7265 |

Gray highlighted cells in each column indicate pairs with a significant difference as assessed using the Nemenyi test for multiple pairwise
comparisons between the groups, P-value < 0.05

Although we demonstrated this capability in the yeast *S. cerevisiae*, we anticipate the approach will be applicable to generating protein level oscillations in many other cell types, considering the ODC degron is functional in vertebrates (Murakami, Matsufuji et al. 1992, Takeuchi, Chen et al. 2008) and non-vertebrate species (DeScenzo and Minocha 1993, Hoyt, Zhang et al. 2003). Light-sensitive degradation using the psd has been successfully demonstrated in cell types including human, and other mammalian cells (Sun, Zhang et al. 2017, Baaske, Gonschorek et al. 2018). Moreover, light-activation is highly suitable for feedback control since the intensity, frequency, and location of illumination can be instantly modulated. Since we demonstrated that controlling the degradation rate is sufficient to drive protein levels under constant transcription, we anticipate this approach to allow generating protein oscillations decoupled from transcriptional signals. This approach may help elucidate the role of oscillations observed in proteins with complex regulatory inputs such as the p53 (Lahav 2004, Porter, Fisher et al. 2016, Harton, Koh et al. 2019) on its downstream gene activation.

## Conclusions

Here we demonstrated robust generation of controlled protein level oscillations in the yeast *S. cerevisiae* using a photo-sensitive degron. We found that by monitoring the total fluorescence of microcolonies of cells, protein oscillations on minute timescales could be reproducibly generated. Using two red fluorescent proteins with distinct maturation times, we showed that the apparent delay in functional protein formation does not change the protein level oscillation generated by modulating the protein degradation rate. The results show that controlling protein degradation rates under constant transcription as a means of modulating protein levels demonstrates a clear pathway for feedback control of protein levels on minute timescales.

## Materials and Methods

### Cloning and expression of a photo-sensitive degron in yeast

The photo-sensitive degron (psd) module used in this study has been previously described by Renicke and colleagues (Renicke, Schuster et al. 2013). A red fluorescent reporter protein (tagRFP) was fused to the N-terminus of *At*LOV2. The tagRFP-psd construct was cloned into the pRS316 plasmid (Huang and Shusta 2005) (a gift from Dr. Eric Shusta) using restriction enzymes NotI and BsrGI (New England Biolabs) under the galactose inducible GAL_1-10_ promoter. A psd without the ODC degron (Δpsd) was generated using polymerase chain reaction and cloned using the same restriction sites. The resulting plasmids were named pRS316-tagRFP-psd and pRS316-tagRFP-Δpsd respectively. The plasmids were transformed into the yeast *Saccharomyces cerevisiae* strain BJ5464 (BJα, α *ura3-52 trp1 leu2*Δ*1 his3*Δ*200 pep4:: HIS3 prb1*Δ*1*.*6R can1 GAL*) cells, using a commercially available kit (Zymo Research). Transformed cells were grown in SD-CAA medium (0.1 M sodium phosphate, pH 6.0, 6.7 g/L yeast nitrogen base, 5.0 g/L casamino acids, 20.0 g/L glucose), supplemented with tryptophan (0.04 g/L) at 30 °C on an orbital shaker (250 rpm).

### qPCR gene expression analysis

Briefly, cells containing pRS316-tagRFP-Δpsd plasmid were grown in SD-CAA medium supplemented with tryptophan (0.04 g/L) at 30 °C on an orbital shaker (250 rpm) overnight. Yeast cells grown overnight were diluted to an OD_600_ of 1, and the medium was replaced with the induction medium, SG-CAA (0.1 M sodium phosphate, pH 6.0, 6.7 g/L yeast nitrogen base, 5.0 g/L casamino acids, 20.0 g/L galactose) supplemented with tryptophan (0.04 g/L) and incubated at 16 °C on an orbital shaker (250 rpm) for 20 hr. The cells were then diluted 5-fold in SG-CAA (supplemented with tryptophan) and grown at 30 °C on an orbital shaker (250 rpm) for period of 400 min with blue light illumination with period of 1/*f* = 100 min. Cells (10 mL of culture) were collected at the end of each 100 min illumination cycle (blue light power density of 0.027 mW/mm^2^ (106.02 µmol/m^2^s)), and stored at 4 °C until analysis. RNA extraction was performed immediately after sample collection using the RNeasy Mini Kit (Qiagen cat# 74104) according to the manufacturer’s protocols. Quality of purified RNAs was checked by measuring the absorbance ratio at 260 nm to 280 nm (A260/A280) by using the NanoDrop ND-1000 (Nanodrop Technologies). The RNA integrity and concentration were assessed by using the automated electrophoresis station Agilent TapeStation 4200 (Bio-Rad).

All RNA samples were diluted in RNase/DNase free water (Fisher Scientific cat# BP28191)) at the same concentration and a volume corresponding to 500 ng was added to iScript^(tm)^ cDNA synthesis kit reagents (Bio-Rad cat# 1708890) for cDNA synthesis, as per the kit instructions. The reaction mixture was incubated for 5 min at 25 °C, followed by 30 min at 46 °C and 5 min at 95 °C and held at 4 °C. The cDNA samples were then stored at –80 °C.

The qPCR was performed by using the SsoAdvanced™ Universal SYBR Green Supermix (Bio-Rad cat# 1725270). 2 μl of cDNA was used per total of 20 μl reaction mixture as a template. *TFC1* and *UBC6* genes were used as housekeeping genes. See Supplementary Fig. 1 and Table 1 for RNA quality control and primers used. Cycling conditions included a polymerase activation step of 98°C for 2 min and 40 cycles at 98°C for 2 sec and annealing /extension at 60°C for 5 sec followed by melt curve analysis from 65 to 95°C in 0.5°C increments. The sizes of amplicons for each gene were verified on a 2% agarose gel. The ΔCt values were calculated by subtracting the threshold cycle (Ct) of *tagRFP* from the average of the Ct of the two reference genes.

### Light-induced degradation and imaging

Yeast cells grown overnight were diluted to an OD_600_ of 1, and the medium was replaced with the induction medium, SG-CAA (0.1 M sodium phosphate, pH 6.0, 6.7 g/L yeast nitrogen base, 5.0 g/L casamino acids, 20.0 g/L galactose) supplemented with tryptophan (0.04 g/L) at 16 °C on an orbital shaker (250 rpm) for 20 hrs. Cells were kept in the dark during the entire induction period to prevent unwanted degradation. Degradation was activated by delivering blue light (470 nm) at a power density of 0.027 mW/mm^2^ (106.02 µmol/m^2^s) using a light-emitting diode (LED, Thorlabs). For imaging tagRFP, green light (530 nm) was delivered at a power density of 0.69 mW/mm^2^.

Additional precautions were taken to prevent exposure to light after induction until the experiment was completed. For imaging, cells were placed in 96-wells plates coated with concanavalin A (2 mg/mL, Sigma Aldrich) to attach the cells to the bottom of the plates. Unbound cells were washed with SG-CAA medium.

The sum of the fluorescence intensity of each microcolony was measured over the experiment to quantify the protein level. To identify regions with microcolonies, ‘Otsu dark’ auto threshold method in the ImageJ library was used. The mean fluorescence intensity over all microcolonies of all five experiments was defined as the normalization factor. The sum for each microcolony was then normalized to the normalization factor. Additionally, an offset specific to each experiment was used in all experimental datasets so that the first value started at a normalized value of 1. In each experiment, measurements from at least 10 microcolonies were recorded and averaged.

LED intensities were controlled with an Arduino (See **Supplementary Information**) and controlled using the µManager (Edelstein, Tsuchida et al. 2014). The system snaps images from all samples every minute while exposing psd and Δpsd samples to a blue LED controlled via the Java programming language (See **Supplementary Information**). The image processing/analysis was conducted using the ImageJ/µManager libraries (See **Supplementary Information** for more details and the codes are available in GitHub (Mahrou 2021)).

### Statistical analysis

To generate the Bode plot, the median phase and amplitude values from each of the five frequencies from ten microcolonies and median of four parameter values estimated for the model were calculated. The median values between the three datasets (Model, tagRFP-psd, and mCherry-psd) were compared using the Nemenyi test. Statistical analyses were performed using real statistics in Excel (Charles Zaiontz, www.real-statistics.com).

## Acknowledgements

This work was funded by the University of Connecticut, the Alzheimer’s Association (2019-AARG-NTF-640971), and NIH (1R21NS111358-01A1). We thank Bo Reese and Lu Li of Center for Genome Innovation within the Institute for Systems Genomics at the University of Connecticut for their help in the RNA quality assessment and qPCR analysis. The authors declare that they have no conflicts of interest with the contents of this article.

## Author Contributions

B.M. designed and performed the experiments, derived the model and analyzed the data. A.P. cloned the psd constructs and conducted the qPCR gene expression analysis. M.H. conducted the statistical analysis. Y.C. and Y.S. supervised the project and assisted with the analysis and the interpretation of data and model. All authors discussed the results and contributed to the manuscript.

