## Supplementary Materials for "Degradation-driven protein level oscillation in the yeast *Saccharomyces cerevisiae*"

**Supplementary Text**

**Cross correlation**

The time delay between input (blue LED light intensity) and output (protein fluorescence intensity) signals can be found by calculating the cross-correlation. The cross-correlation equation reads:

$$G_{u_{\deg}*y}(\tau)=\frac{1}{N-\left| \tau\right|} \sum_{n=0}^{N-\tau+1} u_{\deg}(n)y(n+\tau) \tau\geq0$$

$$G_{u_{\deg}*y}(\tau)=G_{y*u_{\deg}}(-\tau) \tau<0$$

where G_udeg*y_(τ) is the cross-correlation between u_deg_ (input) and y (output) at time lag τ. The time lag τ that corresponds the time delay ($\tau_{\mathrm{delay}})$ is calculated from the minimum value of the G_udeg*y_(τ):

$$\tau_{\mathrm{delay}}\leftarrow\min_{n} \left( G_{u_{\deg}*y}(\tau) \right)$$

Our inputs are in the form of squared waves where outputs are triangular (Fig. 2c). Regardless of the cross-correlation plot acquired from experimental input and output, we can evaluate cross-correlation between schematic squared and triangular waveforms (Supp. Fig. 3). The minimum in the cross-correlation plot occurs at the time lag where the maximum of the input waveform is aligned with the minimum of the output waveform which is equal to the one fourth of the oscillation period (Supp. Fig. 3). We expected to observe the same value for the first minimum which occurs in the cross-correlation plot obtained from the experiment (Fig. 3). However, the minima (indicated by arrows in Fig. 3) are not at the expected times and there are differences between the expected and observed values. The difference is the resulting time delay between the inputs and outputs of the experiments. For example, the minimum of the cross-correlation plot for the experiment with period oscillations of 100 minutes should be at the time lag of 25 minutes (Supp. Fig. 3) but it was observed at 40 minutes. Therefore, the time delay is 15 minutes (40 – 25 minutes). The source code to create experimental cross-correlation is a MATLAB code named CrossCorrelation.m, available on GitHub (Mahrou 2021).

**Normalized least mean square algorithm**

To estimate the parameters of the tagRFP-psd protein turnover system (p_y_, p_pro_, p_deg_) we need to define the estimated system equation from the system model (main text Eq.2):

$$\hat{y}\left( n \right)=\hat{p}_{y}(n-1)y\left( n-1 \right)+\hat{p}_{\mathrm{pro}}(n-1)+ \hat{p}_{\deg}(n-1)u_{\deg}(n-1)$$

where n is the discrete-time index, $\hat{y}$(n) is estimated normalized intensity of tagRFP-psd fluorescence at time n, u_deg_(n) is the normalized intensity of blue light (degradation factor) at time n, and $\hat{p}_{y}$, $\hat{p}_{\mathrm{pro}}$, and $\hat{p}_{\deg}$ are estimated parameters of endogenous degradation, production, and light-induced degradation, respectively. The equation can be re-expressed in a matrix form as follows:

$$\hat{y}\left( n \right)={\hat{\mathbf{p}}\left( n \right)}^{T}\mathbf{u}(n)$$

where $\hat{\mathbf{p}}(n)$ and $\mathbf{u}(n)$ are parameters and inputs vectors, respectively:

$$\hat{\mathbf{p}}(n)=\left[ \begin{matrix} \hat{p}_{y}(n-1) \\ \hat{p}_{\mathrm{pro}}(n-1) \\ \hat{p}_{\deg}(n-1) \end{matrix} \right], \mathbf{u}(n)=\left[ \begin{matrix} y\left( n-1 \right) \\ 1 \\ u_{\deg}(n-1) \end{matrix} \right]$$

The Least Mean Square (LMS) algorithm attempts to minimize squared error e^2^, the error between the measured intensity of tagRFP-psd (or mCherry-psd), y(n), and the estimated intensity of tagRFP-psd (or mCherry-psd), $\hat{y}$(n):

$$e^{2}\left( n \right)=\left( y\left( n \right)-\hat{y}\left( n \right) \right)^{2}=\left( y(n)-{\hat{\mathbf{p}}\left( n \right)}^{T}\mathbf{u}(n) \right)^{2}$$

The parameters in LMS are updated from previous step using steepest descent optimization method as the following:

$$\hat{\mathbf{p}}(n+1)=\hat{\mathbf{p}}(n)-\mu\left( n \right)e\left( n \right)\mathbf{u}(n)$$

Where µ(n) is a chosen step size and can be considered as a constant value. In Normalized LMS (NLMS) µ(n) is self-adjustable and chosen as:

$$\mu\left( n \right)= \frac{\mu}{\left\| \mathbf{u}\mathbf{(}n\mathbf{)} \right\|^{2}}$$

in which µ has a fixed value (0.1 was used for the simulation). The NLMS algorithm follows the below list and the source code is written in MATLAB named NLMS.m and is available on GitHub (Mahrou 2021):

1- Initialize the parameter vector, $\hat{\mathbf{p}}(n)$;
2- Estimate the amount of tagRFP-psd, $\hat{y}$(n);
3- Compute the estimation error, e(n);
4- Update parameters $\hat{\mathbf{p}}(n+1)$;
5- Repeat steps 2 to 4.

After sufficient iterations, the parameters tend to converge. For every experiment, a set of parameters ($\hat{p}_{y}$, $\hat{p}_{\mathrm{pro}}$, and $\hat{p}_{\deg}$) is defined and used to produce the model bode diagram via ModelBode.m MATLAB code (the code is available on GitHub (Mahrou 2021)). Supp. Fig. 5 shows an example of the parameter values resulted from the NLMS estimation.

**Image and data acquisition**

Image and data acquisition are done in two steps. First, the samples are irradiated and the images are saved in a file. Second, the saved images are processed and the fluorescence intensity measurement data are also saved for later analysis. The codes are available in two Java packages named ImageAcquisition and DataAcquisition on GitHub (Mahrou 2021).

The ImageAcquisition package contains Main.java class and two additional classes named InitializeMM.java and ImageSaver.java. InitializeMM.java class loads the Micromanager Configuration and libraries. In addition, user needs to insert the values to set up the stage position for all samples (psd and Δpsd). ImageSaver.java class saves all images in tiff format. Main.java class communicates with the microscope to irradiate samples by blue LED light in a periodic manner and to collect images of all samples every minute. Blue LED light is connected to an Arduino serial port and its operation is programmed using the µManager Java library. The Arduino code is available in the same package named LEDconnect.ino.

DataAcquisition package contains Main.java class and WriteToExcel.java class. The Main.java class reads every image during the iteration, crops the desired cell, extracts it from the background using a threshold method, and quantifies its average fluorescence intensity.

| **Name** | **Sequence** |
| --- | --- |
| P1_F  P1_R | ATGGAAGGCACAGTGAATAACC  GGTTTATGAATGTTCTGGAGCC |
| P2_F  P2_R | ACGGGGGTGTTTTAACTGC  CCTCCTACAAGTTTCAATGCC |
| *TFC1_*F  *TFC1_*R | GCTGGCACTCATATCTTATCGTTTCACAATGG  GAACCTGCTGTCAATACCGCCTGGAG |
| *UBC6_*F  *UBC6_*R | GATACTTGGAATCCTGGCTGGTCTGTCTC  AAAGGGTCTTCTGTTTCATCACCTGTATTTGC |

**Supplementary Table 1**. Primers used in qPCR experiments. Two different primer pairs were used to amplify the *tagRFP* gene. *TFC1* and *UBC6* were used as reference housekeeping genes due to their previously validated stable expression in *S. cerevisiae*.

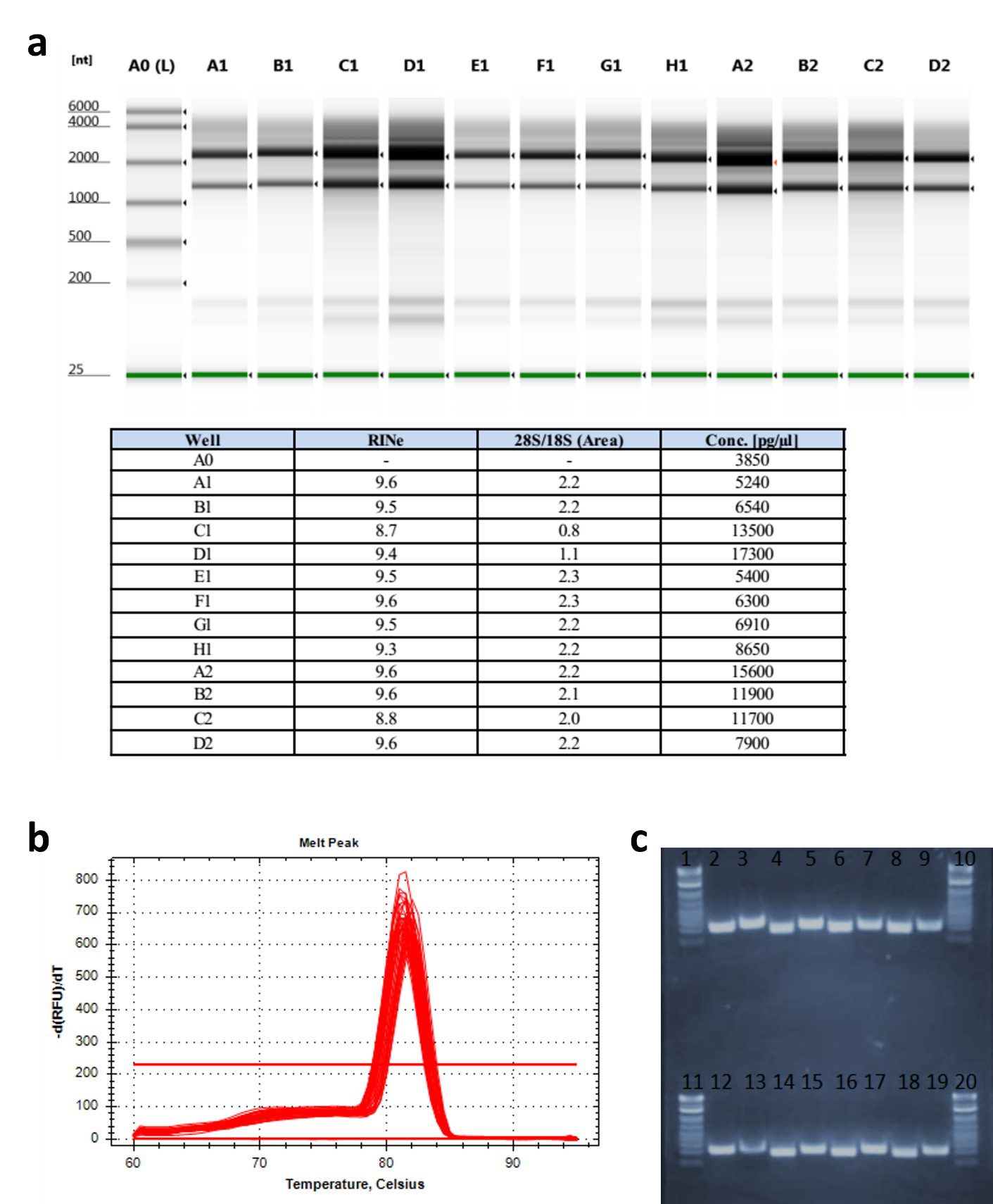

**Supplementary Figure 1. Quality assessment of RNA samples used for qPCR.** (**a**) Gel electrophoresis of yeast extracted RNA using the Agilent Tapestation 4200. The associated RNA Integrity Number (RINe), 28S/18S rRNA ratio, and concentration are shown below the gel images. Lanes A1-D1, E1-H1 and A2-D2 represents RNA samples from 3 different experiments, respectively (each from 0, 100, 200, and 300 minutes time points). (**b**) The melting curves obtained from the qPCR. (**c**) Aliquots of qPCR products were analyzed on a 2% agarose gel to determine amplicon molecular weight and amplification specificity: lanes: 1, 10, 11,20 – Ladder; 2, 12 – *TFC1* at time 0 and 300 minutes ; 3, 13 – *UBC6* at time 0 and 300 minutes; 4, 14 – *P1* at time 0 and 300 minutes (n = 1); 5, 15 – *P2* at time 0 and 300 minutes (n = 1); 6, 16 – *P1* at time 0 and 300 minutes (n = 2); 7, 17 – *P2* at time 0 and 300 minutes (n = 2); 8, 18 – *P1* at time 0 and 300 minutes (n=3); 9, 19 – *P2* at time 0 and 300 minutes (n = 3). P1 and P2 represent two different primer pairs for the *tagRFP* gene. The primer sequences are provided in Supp. Table 1.

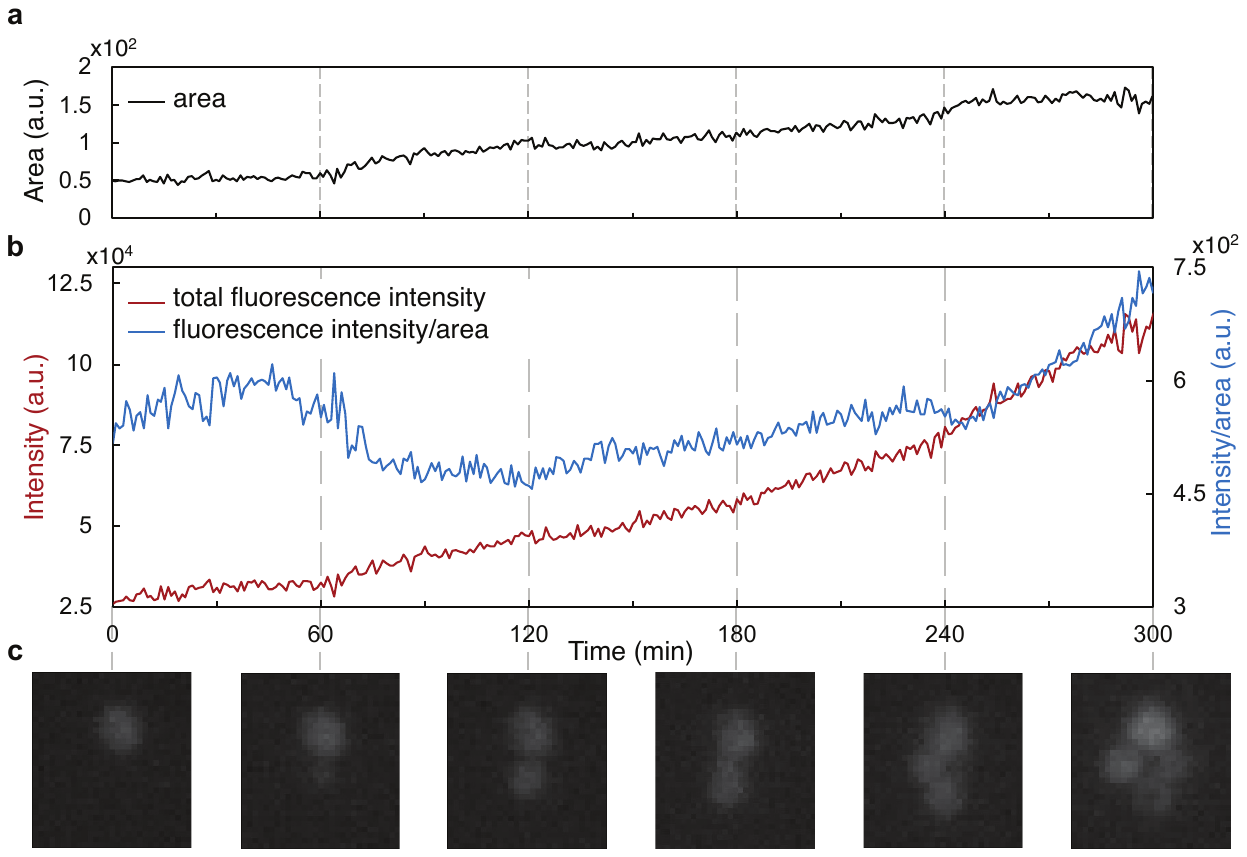

**Supplementary Figure 2. The effect of cell division on fluorescence intensity.** (**a**) Area (number of pixels as determined using ‘Otsu dark’ auto threshold in ImageJ library) of the target cell and its newly generated daughter cells measured over time. (**b**) Blue and red plots indicate fluorescence protein intensity density (the sum of fluorescence intensity over the cell area) and the total fluorescence intensity (summing the fluorescence intensity of all pixels), respectively. (**c**) Microscope images of yeast cells corresponding to the indicated timepoints in panel (b).

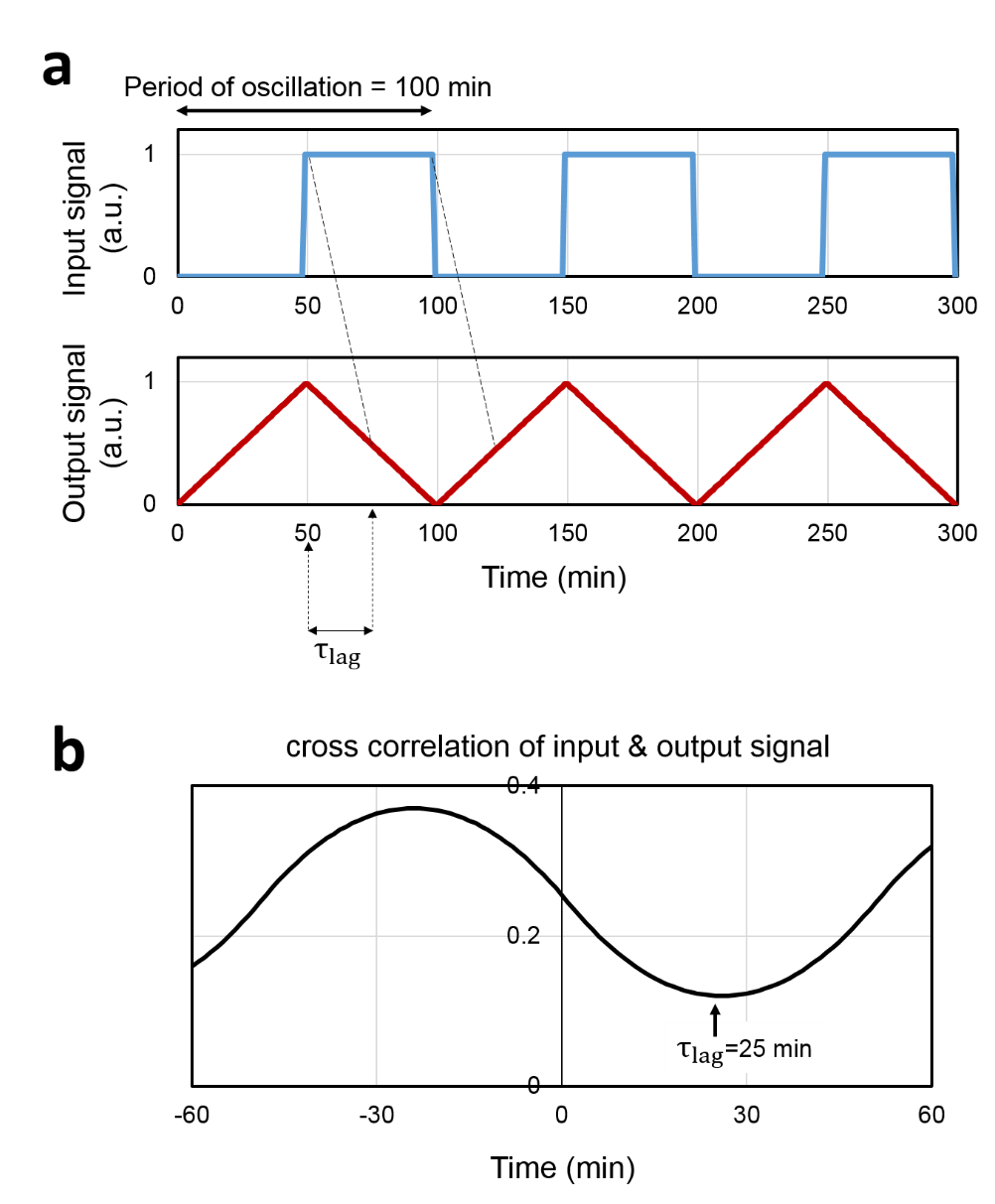

**Supplementary Figure 3. Determination of the minimum cross-correlation between squared and triangular waveforms.** (**a**) Hypothetical input and output waveforms with cycle period of 100 minutes. The τ_lag_ is the time delay corresponding to minimum cross-correlation. The minimum occurs at a time lag equal to one fourth of the oscillation period. This is where the maximum of the input waveform is aligned with the minimum of the output waveform. (**b**) Shows the cross-correlation of the two input and output signals. The minimum of the plot occurs at a time lag of 25 minutes, that is one fourth of the period of the oscillations (100 minutes).

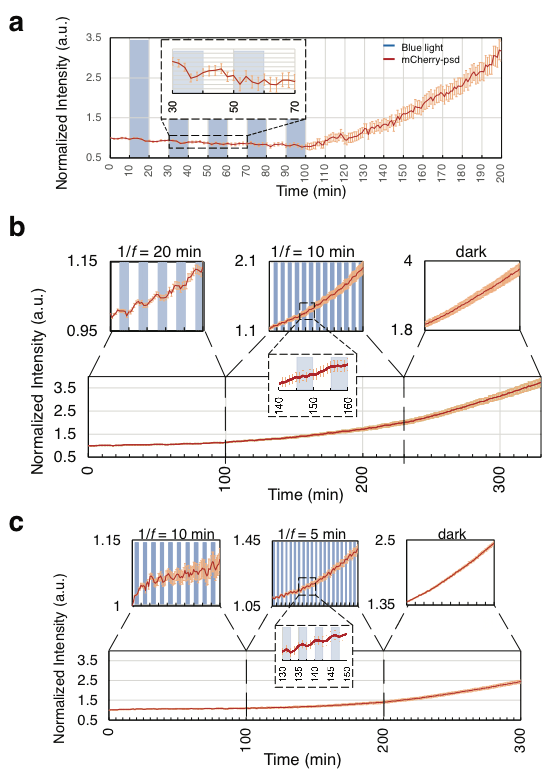

**Supplementary Figure 4. Extended experimental results for protein turnover in the cells producing mCherry-psd.** (**a**) mCherry-psd under varying frequencies of blue light illumination. Input illumination frequency was 1/*f* = 20 minutes during the first 100 minutes, after which the blue light was off. (**b**) mCherry-psd fluorescence oscillations under input frequency 1/*f* = 20 minutes for 100 minutes, followed by 1/*f* = 10 minutes for 130 minutes, then followed by no blue light for 100 minutes. (**c**) mCherry-psd fluorescence oscillations under input frequency 1/*f* = 10 minutes for 100 minutes, followed by 1/*f* = 5 minutes for 100 minutes, then followed by no blue light for 100 minutes. For all panels, solid lines indicate the mean fluorescence intensities and lighter lines indicate standard deviation from n = 25 to 30 microcolonies. Blue light intensity was 0.027 mW/mm^2^ (106.02 µmol/m^2^s)) for all panels.

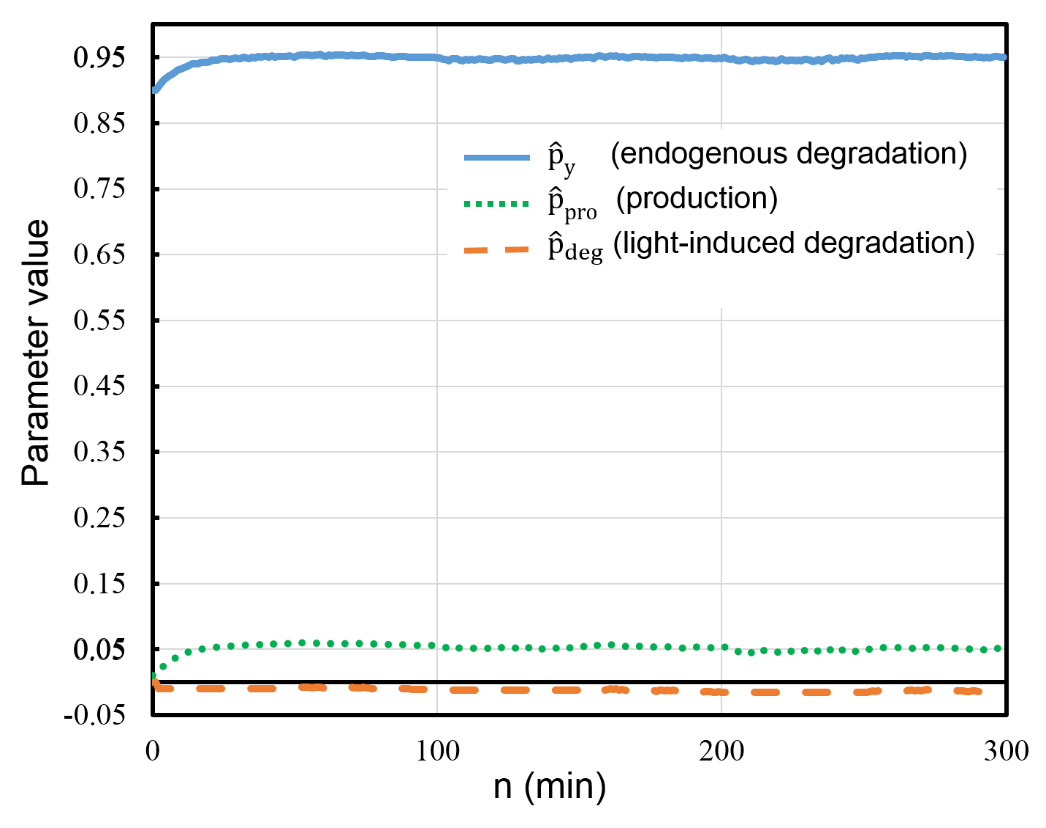

**Supplementary Figure 5. Parameter estimation.** A representative NLMS parameter estimation for the tagRFP-psd related to the experiment under input illumination frequency of 1/*f* = 100 minutes. The parameters converge to values of $\hat{p}_{y}$≈ 0.95, $\hat{p}_{\mathrm{pro}}$≈ 0.05, and $\hat{p}_{\deg}$≈ -0.01.
